# Thermogenic Adipose ADH5 Counteracts Age-related Metabolic Decline

**DOI:** 10.1101/2025.07.01.662628

**Authors:** Sara C. Sebag, Tate Neff, Qingwen Qian, Arvand Asghari, Zhuozhi Wang, Zeyuan Zhang, Mark Li, Meihua Hao, Vitor A. Lira, Hongli Sun, Matthew J. Potthoff, Ling Yang

**Author notes:** Equal contribution (Sara C. Sebag, Tate Neff). Corresponding author: Address all correspondence to: Ling Yang, Ph.D., Department of Anatomy and Cell Biology, Fraternal Order of Eagles Diabetes Research Center Pappajohn Biomedical Institute, University of Iowa Carver College of Medicine, Iowa City, IA 52242.

## Abstract

Aging-associated decline in brown adipose tissue (BAT) function and mass contributes to energy and metabolic homeostasis disruption. Alcohol dehydrogenase 5 (ADH5) is a major denitrosylase that prevents cellular nitro-thiol redox imbalance, an essential feature of aging. However, the functional significance of BAT ADH5 in the context of aging is largely unknown. Here, we aimed to investigate the role of BAT ADH5 in protecting against age-related metabolic dysfunction. We show that aging elevates BAT protein S-nitrosylation modification and downregulates ADH5 in mice. Furthermore, BAT ADH5-deletion accelerates BAT senescence and aging-associated declines in metabolic homeostasis and cognition. Mechanistically, we found that aging inactivates BAT *Adh5* by suppressing heat shock factor 1 (HSF1), a well-recognized proteostasis regulator. Moreover, pharmacologically enhancing HSF1 improved age-related BAT senescence, metabolic decline, and cognitive dysfunction. Together, these findings suggest that the BAT HSF1-ADH5 signaling cascade plays a key role in protecting against age-related systemic functional decline. Ultimately, unraveling the role of thermogenic adipose nitrosative signaling will provide novel insights into the interplay between BAT nitric oxide bioactivity and metabolism in the context of aging.

**Highlights:**

- Aging elevates general protein S-nitrosylation while downregulating ADH5 expression in BAT.
- Loss of thermogenic adipose tissue ADH5 accelerates BAT senescence and exacerbates aging-associated metabolic and cognitive dysfunctions.
- Disruption of HSF1-activated *Adh5* in BAT contributes to aging-associated metabolic and cognitive impairments.
- Pharmacological targeting of BAT HSF1 alleviates aging-related BAT senescent and systemic declines.

## 1. INTRODUCTION

Brown adipose tissue (BAT) is essential for classical nonshivering thermogenesis by utilizing intracellular and systemic lipid and glucose fuels[1–3]. Accumulated evidence shows longer-lived mice have highly active BAT presenting increased mass and thermogenic-metabolic function[4; 5]. In humans, the beneficial effects of BAT depots have been demonstrated against deleterious outcomes of aging and age-related diseases such as cardiovascular disease (CVD)[6]. Notably, in rodents and humans, improvement of BAT activity ameliorates aging-mediated hyperglycemia and hyperlipidemia[7–12]. However, BAT mass and function decline with age leading to impaired BAT metabolic capacity, the onset of age-related diseases and, ultimately, shortened lifespan[13–16]. Therefore, it is urgent to understand the multifaceted cellular and systemic factors that promote age-associated pathologies to design new therapeutics for healthy aging.

The mechanisms underlying aging are diverse and complex involving cell autonomous as well as systemic and environmental factors. These include increased production of reactive oxygen and nitrogen species (ROS and RNS, respectively) paired with reduced antioxidant capacity, mitochondrial defects[17–19], cellular senescence, and metabolic dysfunction[19–23]. Moreover, aged and senescent cells accumulate damaged proteins leading to disrupted proteostasis[24]. Although mitochondrial dysfunction, aberrant redox network remodeling, and proteostasis are key features in aging as well as deleterious factors implicated in BAT dysfunction[25–31], the molecular mechanisms and modulators linking these signaling cascades in BAT aging are largely unknown.

Intracellularly, ROS and RNS are redox mediators that regulate diverse aspects of cellular processes including modulating protein function[32]. For example, redox modification of protein cysteine residues plays critical roles in physiology and pathologies including aging[32]. In BAT, it has been demonstrated that acute thermogenic induction generates physiologic levels of mitochondrial ROS which activates uncoupling protein 1 (UCP1) by sulfenylation[28; 33]. Furthermore, cysteine oxidation networks in BAT are selectively modified in young mice and lost in aged animals[34]. In contrast, Kazak et al. reported that UCP1 deficiency elevated BAT ROS[35], implicating a bidirectional interaction between ROS and UCP1. Excessive nitric oxide (NO) production and the resultant aberrant NO-mediated protein cysteine nitrosylation (S-nitrosylation, SNO) induce nitrosative stress which disrupts essential aspects of cellular function including organelle homeostasis, inflammatory responses, and metabolism, all of which decline during aging[36; 37]. However, the functional significance of RNS in aging-related BAT dysfunction remains uncharacterized.

Alcohol dehydrogenase 5 (ADH5; also named S-nitrosoglutathione reductase, GSNOR) is the major cellular denitrosylase that regulates NO availability in the cell by catalyzing the breakdown of SNOs and balancing the intracellular thiol redox status[36; 38; 39]. Recently we demonstrated that BAT ADH5 protects against obesity-associated metabolic dysfunction[40]. Here we revealed the functional impact of the proteostasis master regulator, heat shock factor 1 (HSF1), on BAT ADH5 in the context of aging. Our study provides a novel insight into the BAT NO bioactivity-linked interplay between proteostasis and metabolic dysfunction in aging.

## 2. MATERIALS AND METHODS

### 2.1 Nanoclay-mediated drug delivery

Nanoclay (NC, Laponite XLG) was obtained from BYK Additives Inc. TX, USA. Rat tail Type I collagen (RatCol) and Neutralization solution were procured from Advanced Biomatrix, Carlsbad, CA (5153). HSF1A was bought from Millipore, Burlington, MA (1196723-93-9). For the Collagen-NC gel, 850 µL of cold collagen stock solution (4.1 mg/ml) was added to 50 µL of Nanoclay stock solution (10 mg/ml). For collagen-NC-HSF1A, 50 µL of Nanoclay stock solution (10 mg/ml) was mixed with 3.33 µL HSF1A and then mixed with 850 µL of cold collagen stock solution (4.1 mg/ml). All gels were supplemented with 90 µL neutralization buffer and kept on ice prior to sub-BAT injection.

Based on the structure of HSF1A, the concentration was determined by UV-Vis spectroscopy. Firstly, a standard curve of HSF1A was obtained by measuring the UV-Vis spectrum of different concentrations of HSF1A in PBS (1.25, 2.5, 5, 10, 20 µM). Thereby, two standard curves at 207 nm and 284 nm were generated. Next, two types of collagen gels with or without Nanoclay were fabricated to study the release profile. Briefly, 50 µL of DI water (for Col-HSF1A) or Nanoclay stock solution (for Col-NC-HSF1A) was mixed with 3.33 µL of HSF1A and the mixture was added to 850uL of cold collagen solution and mixed well with a pipette. Then 90 µL of neutralization buffer was added and mixed with a pipette. Finally, the mixed solution was divided into 3 parts and placed at 37°C for 30 min. After gelation, 1mL of fresh PBS buffer was added to each tube for HSF1A release. At each time point (1, 5, 9, 14, 21 days), the PBS buffer in each tube was collected for UV-Vis spectroscopy.

### 2.2 Mouse models

Animal care and experimental procedures were performed with approval from the University of Iowa’s Institutional Animal Care and Use Committee. Animals received humane care in compliance with the *Guide for the Care and Use of Laboratory Animals* (National Academies Press, 2011) and with the Principles of Laboratory Animal Care formulated by the National Society for Medical Research. C57BL/6J mice (The Jackson Laboratory, 000664), UCP1-cre (The Jackson Laboratory, B6.FVB-Tg(UCP1-cre)1Evdr/J, 024670), and *Adh5*^fl^ [41] mice were kept on a 12-hr light/dark cycle. At 9-11 months, NC-Collagen or NC-HSF1A was interscapularly administrated at a dosage of 1 mM/mouse or 0.2 mg/kg. For indirect calorimetry analysis, 12-month-old mice were singly housed in an environmental chamber during a 12-hour light cycle (Metabowl; Jencons Scientific). Mice were given access to standard 2920X Teklad Global Diet and water *ad libitum* while in cages. All tissues were harvested, frozen in liquid nitrogen, and kept at −80°C until processed.

### 2.3 BAT explants treatment

For HSF1 inhibition, BAT explants from WT mice were exposed to 2 µM KRIBB11 (Sigma, 385770) for 24 hr. For HSF1 activation, explants were treated for 24 hr with HSF1A (20 µM). All tissues were harvested, frozen in liquid nitrogen, and kept at −80°C until processed.

### 2.4 Quantitative Real-time RT-PCR

Total RNA was isolated using TRIzol reagent (Invitrogen,15-596-018) and reverse transcribed into cDNA using the iScript cDNA synthesis kit (Bio-Rad, 1708890). Quantitative real-time RT-PCR analysis was performed using SYBR Green (Invitrogen, KCQS00). Primer sequences are found in Table S1.

### 2.5 Chromatin immunoprecipitation (ChIP) assay

The chromatin immunoprecipitation assay was performed using the SimpleChIP Enzymatic Chromatin IP Kit (Cell Signaling Technology, 9003) with some modifications. Briefly, BAT was crosslinked with 1% formaldehyde (Sigma-Aldrich, F8775), after which the reaction was stopped by washing with ice-cold PBS (Gibco, 14,190,144) containing 0.125 M glycine (americanBio, AB00730-10,000) and protease inhibitor (Sigma-Aldrich, P8849). Nuclei were then isolated, and the chromatin was immunoprecipitated with protein A/G magnetic beads (Thermo Fisher Scientific, 88,802) conjugated with anti-IgG (Cell Signaling Technology, 2729) and anti-HSF1 (Cell Signaling, D3L8I, 12972) overnight at 4°C. The DNA was eluted from the beads and subjected to PCR analysis. The primers used for ChIP assays were mouse *Adh5* Forward: -1735 to -1716; TGCTCCTACTCTCTCTCCCC; mouse *Adh5* Reverse: -1645 to -1624; AGCCCTTTTGTTCCTTTTCACA.

### 2.6 Western blot analysis

Proteins were extracted from BAT and subjected to SDS–polyacrylamide gel electrophoresis, as previously described[42]. Membranes were incubated with anti-CoxI (Santa Cruz, H-1, sc-166573), anti-CoxII (Santa Cruz, H-3, sc-376861), anti-CoxIII (Santa Cruz, N-20, sc-23986), anti-CoxIV (Santa Cruz, C-20, sc-23982) or anti-Cox5a (Santa Cruz,(A-5), sc-376907) antibodies at 1:1000 dilution and then incubated with the appropriate secondary antibody conjugated with horseradish peroxidase (1:5000, Santa Cruz, sc-2005 or 1:5000, Cell Signaling Technology, 7074S). The signal was detected using the ChemiDoc Touch Imaging System (Bio-Rad), and densitometry analyses of western blot images were performed by using the Image Lab software (Bio-Rad).

### 2.7 S-nitrosylation detection

#### In situ detection of S-nitrosylated protein

The assay was performed as described with minor modifications[43]. Briefly, frozen BAT sections were fixed with 3.5% paraformaldehyde, and then permeabilized with 0.1% Triton X-100 in PBS containing 1 mM EDTA and 0.1mM neocuproine. Biotin-switch assay was performed by first blocking free thiols using HENS buffer containing 20 mM MMTS at room temperature for 30 min. Then the S-nitrosylated proteins were labeled in HENS buffer with 0.4 mM biotin-HPDP and 10 mM ascorbic acid for 1-hr. Biotinylated proteins were labeled using streptavidin conjugated with Alexa 488 (ThermoFisher Scientific, A32731). The images were observed using a Zeiss 700 confocal or Leica fluorescence microscope and quantified using ImarisColoc (Bitplane).

#### Detection of S-nitrosylation proteins

Total S-nitrosylated proteins in BAT were detected by S-nitrosylation Western Blot Kit (Thermo Fisher Scientific). Briefly, BAT was lysed in HEN buffer and 500 µg protein was used for each sample. Free cysteines were first blocked with MMTS for 20 min at 50°C. Following precipitation, using 4× volume of cold acetone for 60 min at −20°C, S-nitrosylated cysteines were selectively labeled with iodoTMT reagent in the presence of 20 mM sodium ascorbate at RT for 2 hr. Samples were then separated by SDS-PAGE and blotted to the membrane for detection of SNO proteins by the anti-TMT antibody.

### 2.8 Immunohistochemistry, Immunofluorescence and ELISA

For immunohistochemistry, iBAT or iWAT depots were fixed with 4% PFA and sectioned at 5 μm thick, followed by deparaffinization and rehydration processes. Tissue sections were stained using H&E. The images were observed under a Nikon microscope (10x). For immunofluorescence (IF), sections were blocked and incubated overnight at 4°C with appropriate antibody including F4/80 (Cell Signaling, D4C8V, 30325), p16 INK4A (Cell Signaling, E6N8P, 18769) and HSF1 (Cell Signaling, D3L8I, 12972) at 1:100 followed by anti-rabbit (Cell Signaling, 7074) for one hour at RT (1:2000). Nuclei were stained with 4′,6-diamidino-2-phenylindole (1:2500, DAPI). Images were taken using an LSM 880 confocal microscope (Carl Zeiss) and analyzed with NIH ImageJ.

For IL 1β measurement, multiplex cytokine analysis was performed via Bio-Rad Laboratories Bio-plex Pro Mouse Cytokine 23-Plex according to the manufacturer’s instructions for plasma cytokine analysis. Samples were analyzed on a Bio-Rad Laboratories Bio-Plex (Luminex 200) analyzer in the University of Iowa Flow Cytometry Core Facility.

### 2.9 Transmission Electron Microscopy (TEM)

Brown adipose tissues were removed and cut into ∼1 mm^3^ cubes. All electron microscopy (EM)-related reagents were from Electron Microscopy Sciences. Samples were fixed in 2.5% glutaraldehyde and 4% formaldehyde cacodylate buffer overnight (16 hours) at 4°C. Tissue was postfixed in fresh 1% OsO_4_ for 1 hr, dehydrated using a graded alcohol series followed by propylene oxide, and embedded in Epon resin as previously described[44]. Resin blocks were trimmed with a glass knife, cut to ultrathin (50–70 nm) sections with a diamond knife, and mounted on Formvar-coated copper grids. Grids were double contrasted with 2% uranyl acetate and then with lead citrate. Images were captured by a Hitachi HT7800 transmission electron microscope. Cell area was determined using ImageJ.

### 2.10 Barnes maze and rotarod studies

Barnes Maze [46] testing was performed on a 91cm diameter round surface with holes of 5cm diameter equally spaced around the edge (Noldus). The escape chamber was placed randomly under one of the holes and remained in place throughout training. The maze was surrounded with white cloth baring four visual markers for orientation, and a light was placed above the maze. Each mouse was trained for four days, with four trials in each day. For mice that did not locate the escape chamber within 3 minutes, they were guided to the hole with the escape chamber. Short and long term memory based on spatial cues was measured using the Ethovision XT 11.5 video tracking software (Noldus) and confirmed by manual tracking. Rotarod testing took place after training on the Rotamex-5 rotarod (Columbus Instruments). The rotarod spun at 5 rpm, time on the rod until completely falling off was measured at regular intervals with a maximum of 60 seconds.

### 2.11 Metabolic Phenotyping

#### Whole-body energy expenditure and body composition and glucose tolerance

Whole-body energy expenditure (VO_2_, VCO_2_), food intake, and locomotor activity were monitored using a Comprehensive Lab Animal Monitoring System (CLAMS, Columbus Instruments) at the Fraternal Order of Eagles Metabolic Phenotypic Core. Body composition was measured by using Bruker Minispecs (LF50). For the Glucose tolerance test, animals were fasted for 16 hr prior to GTT. Glucose tolerance was tested by measuring glucose concentration at different time points after an intraperitoneal (IP) glucose injection (0.8-1.25 g/kg body weight, 50% dextrose, Hospra Inc, 0409-6648-02)[45].

### 2.12 Echocardiography

Transthoracic echocardiograms were performed on conscious mice in the University of Iowa Cardiology Animal Phenotyping Core Laboratory using a VisualSonics Imaging system using a 30 Mhz transducer. The echocardiographer was blinded to the treatments and genetic backgrounds of the animals. Short-axis images were acquired parallel to the mitral valve plane to obtain the largest cross-sectional image of the left ventricle which included papillary muscles. Long-axis views were obtained perpendicular to the mitral valve plane and were deemed optimal when the diastolic apex-to-base length was the longest including the aortic valve with ascending aorta.

### 2.13 Metabolomic Analysis

#### Tissue samples

Whole BAT tissue was homogenized in ice-cold extraction buffer containing 2 parts methanol, 2 parts acetonitrile, 1 part water with internal standards (Cambridge Isotope Laboratories). Samples were then rotated for an hour at -20°C prior to centrifugation (10 minutes, 21,000 x g). 150 µl of the cleared extracts were moved to autosampler vials and vacuum dried (SpeedVac vacuum concentrator, Thermo). Samples were then reconstituted in 30 µl of methoxyamine (MOX, 11.4 mg/ml) in anhydrous pyridine, vortexed for 5 minutes, and heated for an hour at 60°C. 20 µl of N, O-Bis(trimethylsilyl)trifluoroacetamide (TMS) was added to each sample, which were then vortexed for another minute and heated for 30 minutes at 60°C.

#### GC-MS method

Derivatized samples were analyzed by GC-MS. 1 μl of derivatized sample was injected into a Trace 1300 GC (Thermo) fitted with a TraceGold TG-5SilMS column (Thermo) operating under the following conditions: split ratio = 20:1, split flow = 24 μl/min, purge flow = 5 ml/min, carrier mode = Constant Flow, and carrier flow rate = 1.2 ml/min. The GC oven temperature gradient was as follows: 80°C for 3 min, increasing at a rate of 20°C/min to 280°C, and holding at a temperature of 280°C for 8-min. Ion detection was performed by an ISQ 7000 mass spectrometer (Thermo) operated from 3.90 to 21.00 min in EI mode (-70eV) using select ion monitoring (SIM).

#### Data analysis

Raw data were analyzed using TraceFinder 5.1 (Thermo). Metabolite identification and annotation required at least two ions (target + confirming) and a unique retention time that corresponded to the ions and retention time of a reference standard previously determined in-house. A pooled sample generated prior to derivatization was analyzed at the beginning, at a set interval during, and at the end of the analytical run to correct peak intensities using the NOREVA tool[46]. NOREVA corrected data were then normalized to the total signal per sample to control for extraction, derivatization, and/or loading effects.

### 2.14 Quantification and Statistical Analysis

Results are expressed as the mean ± the standard error of the mean (SEM); *n* represents the number of individual mice (biological replicates) or individual experiments (technical replicates) as indicated in the figure legends. We performed the Shapiro-Wilk Normality test in experiments that have a relatively large sample size (n>5) and found that these data pass the normality test (alpha=0.05). Data were further analyzed with two-tailed Student’s and Welch’s *t*-test for two-group comparisons or ANOVA for multiple comparisons. For both One-Way ANOVA and Two-Way ANOVA, Tukey’s post-hoc multiple comparisons were applied as recommended by Prism. In all cases, GraphPad Prism (GraphPad Software Prism 8) was used for the calculations.

## 3. RESULTS

### 3.1 Aging elevates thermogenic adipose protein S-nitrosylation

To investigate the pathophysiological relevance of nitrosative signaling in BAT in the context of aging, we examined the general protein SNO status in BAT from young (3 and 6--month) and old mice (12 and 22-month) male C57BL/6J mice. Using both immunofluorescence (IF) and an iodoTMT-based S-nitrosylation (SNO) analysis, we found increased BAT-SNO of general proteins in older animals compared to young mice, (Fig. 1A&B). Protein S-nitrosylation is regulated by both nitric oxide synthase (NOS)-mediated NO generation and cellular denitrosylation[36] which is tightly controlled by the major denitrosylase, alcohol dehydrogenase 5 (ADH5; also named S-nitrosoglutathione reductase)[36; 39]. Here, we found that aging downregulated *Adh5* in both BAT and inguinal white adipose tissue (iWAT) (Fig. 1C&D), as well as a significant increase in BAT expression of iNOS, a key inducer for protein S-nitrosylation (Fig. 1C).

**Figure 1.**
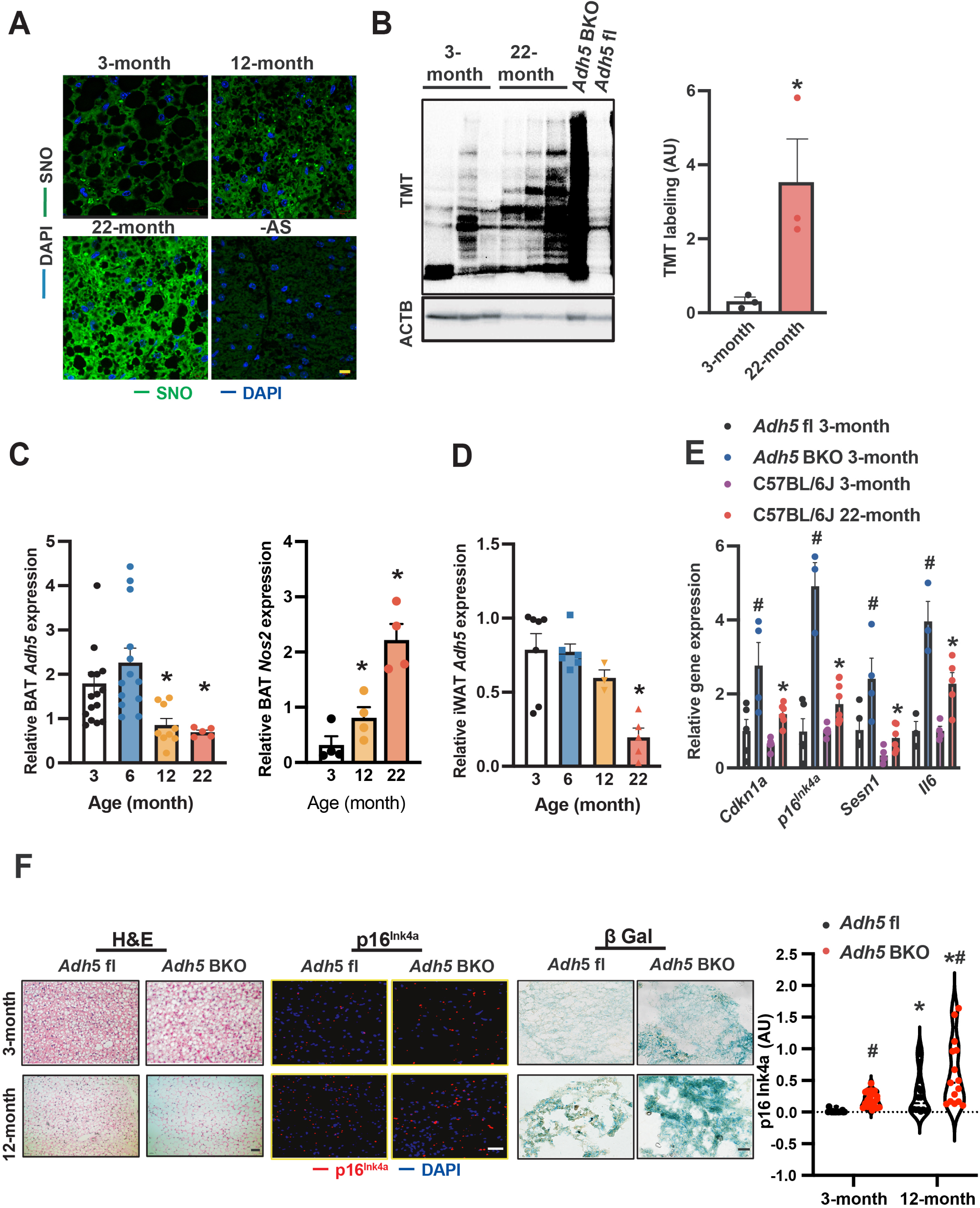
Age-induced BAT ADH5 reduction contributes to BAT senescence. **A.** Representative images (63X) of protein S-nitrosylation staining (SNO) in BAT from young (3-month) or old mice (12- and 22-month). –AS: no ascorbate; negative control for SNO staining. Scale bar: 10 μm. **B.** Representative Western and densitometric analysis (normalized to ACTB input) of SNO proteins in BAT from 3- or 22-month-old wild-type mice. 6-month-old Adh5 BKO and Adh5 fl/fl mice were used as positive and negative control respectively. **C&D.** *Adh5 and Nos2* (encodes iNOS) expression (normalized to *Gapdh*) in BAT and iWAT respectively of *Adh5^fl^ and Adh5*^BKO^ mice. N= 3-7 mice/group. **E.** Levels of mRNAs in BAT of young (3-month) *Adh5^fl^, Adh5*^BKO,^ or WT mice compared to 22-month-old WT by RT-qPCR. N= 3-5 mice/group. **F.** Representative images of H&E, IF of p16^lNK4a^, and β gal staining in BAT from age-matched *Adh5^fl^*and *Adh5*^BKO^ mice. Scale bar: 10 μm in H&E and p16^lNK4a^, 50 μm in β gal staining. Right panel: quantified fluorescence intensities of p16^lNK4a^. 5 fields/mouse, n= 3 mice/group. Data are presented as means ± SEM. *indicates statistical significance relative to the 3-month groups, and ^#^indicates genetic effects in mice at the same age; as determined by Student’s *t*-test in (B), one-way ANOVA in (C&D) and two-way ANOVA in (E&F) followed by posthoc test, p<0.05.

To determine the functional role of BAT-ADH5 in the context of aging, we next generated BAT ADH5 knockout (*Adh5*^BKO^) mice by crossing *Adh5*^fl/fl^ mice (in a C57BL/6J background)[41] with a *Ucp1*-Cre transgenic mouse line[40]. One of the major drivers for the age-related decline is the induction of cellular senescence which is characterized by the induction of cell cycle inhibitors such as *p16^INK4a^* and *Cdkn1a(p21)* as well as senescence-associated secretory phenotype (SASP) markers (e.g., *Il6 and Sesn1*)[31; 47; 48]. As shown in Fig. 1E, BAT-ADH5 deletion increased senescence markers compared with *Adh5*^fl^ mice at 3 months of age. Notably, there were comparable levels of senescent markers in BAT of young *Adh5*^BKO^ mice with aged wild-type mice (Fig. 1E). Consistently, IF analysis and senescence-associated β-galactosidase (β gal) staining revealed increased p16^Ink4a^ in *Adh5*^BKO^ mice compared with its age-matched littermate controls (Fig. 1F). Collectively, these data indicate aging is associated with aberrant BAT ADH5-mediated nitric oxide activity, and loss of AHD5 accelerates BAT senescence.

### 3.2 BAT ADH5 deletion worsens age-associated functional decline

Redistribution of fat depots is a key feature of aging[49; 50]. Consistently, we found an age-associated increase in body weight and that loss of BAT ADH5 accelerated age-associated adiposity and lean mass loss (Fig. 2A&B). BAT is essential for maintaining whole-body energy expenditure and glucose homeostasis[51]. Therefore, we next determined the effect of BAT-ADH5 deletion on systemic energy and metabolic homeostasis in young (3 months) and aged mice (12 months). Metabolic profiling using indirect calorimetry assessment revealed that 12-month-old *Adh5*^BKO^ mice had reduced heat and oxygen consumption compared with age-matched controls (Fig. 2C&D). These systemic energy balance results were further supported by worsened glucose intolerance in both young and older BAT-ADH5 deficient mice (Fig. 2E&F). Of note, the glucose intolerance in 3-month-old *Adh5*^BKO^ mice was comparable to 12- month wildtype mice, indicating a potential for accelerated age-reduced metabolic function in mice with BAT *Adh5* deletion.

**Figure 2.**
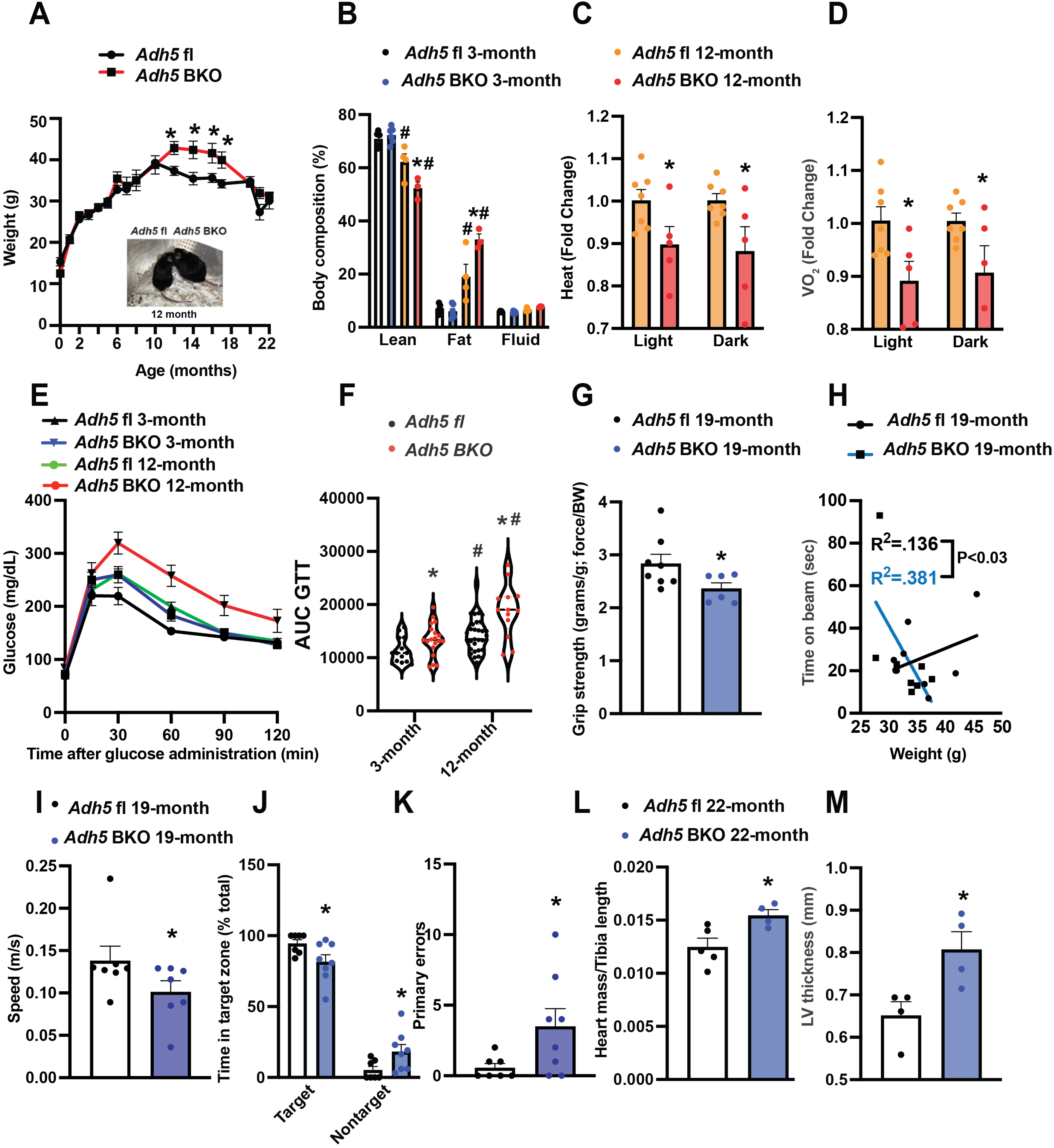
BAT *Adh5* deletion accelerates age-related functional declines. **A.** Body weight (n= 5-22 mice/group), and **B**. Body composition of *Adh5*^fl^ and *Adh5*^BKO^ mice (n= 4-6 mice/group) at indicated age. Inset: representative picture of *Adh5*^fl^ and *Adh5*^BKO^ mice at 12 months. **C&D**. Whole-body energy balance measured in *Adh5*^fl^ and *Adh5*^BKO^ mice housed within a metabolic cage (n= 5-7 mice/group). **E-F**. GTT and AUC from *Adh5*^fl^ and *Adh5*^BKO^ mice (n= 14-30 mice/group). **G.** Grip strength, **H.** Rotarod assessment, **I.** Barnes maze assessment of velocity, **J**. percent of time in target/nontarget zone and **K.** primary errors to escape hole in 19-month-old *Adh5*^fl^ and *Adh5*^BKO^ mice. N = 7-9 mice/group. **L&M.** Heart mass and left ventricle thickness (LV) from age-matched *Adh5^fl^* and *Adh5*^BKO^ mice measured by echocardiography; n = 5-7 mice/group. Data are presented as means ± SEM. *indicates genetic effects in same-age mice, and #indicates age effects in the same type of mice; as determined by Student’s *t*-test (A, C, D, and G-M), and ANOVA in (B) or ANOVA of AUC in (F), p<0.05.

Aging reduces muscle quality[52], strength[53; 54], and lower extremity performance[55]. We next compared sensorimotor changes in aged *Adh5*^BKO^ mice with littermate controls. As shown in Fig. 2G&H, ADH5 deletion in the BAT significantly reduced grip strength and motor performance in 19-month-old mice compared to controls. Aging is a complex process associated with progressive cognitive impairment[56]. To answer whether and to what extent BAT ADH5 deletion alters murine neurological function, we next performed the Barnes maze test[57] in aged *Adh5*^BKO^ mice and its littermate controls. Compared with controls, *Adh5*^BKO^ mice exhibited reduced velocity (Fig. 2I), spatial learning, and short-term memory (decreased frequency within the target zone and increased primary errors) (Fig. 2J&K). Aging is a major risk factor for heart dysfunction whereas BAT protects against age-related diseases such as CVD[6]. Echocardiography revealed increased heart mass and LV thickness in 22-month-old *Adh5*^BKO^ mice compared to age-matched controls (Fig. 2L&M). Together these data suggest that BAT ADH5 is required for maintaining metabolic, motor, cognitive, and cardiac functions in aging.

### 3.3 BAT ADH5 deletion augments aging-related inflammation and mitochondrial defects

Aging is characterized by chronic low-grade systemic inflammation as well as disruption of innate and adaptive immunity[58–62]. We found that BAT-specific *Adh5* deletion increased proinflammatory markers in the SVF of BAT as well as serum IL-1β from young *Adh5*^BKO^ mice compared to *Adh5*^fl^ mice (Fig. 3A&B). To determine the effect of aging on ADH5-regulated BAT inflammation, we next measured F4/80 expression in BAT from aged *Adh5*^BKO^ and *Adh5*^fl^ mice. As shown in Fig. 3C, loss of the BAT-ADH5 increased the infiltration of F4/80-positive cells in both young and aged mice. Beige adipocytes residing in iWAT are activated by β3-adrenergic receptor stimulation leading to browning[63]. We also found an increase in F4/80-positive cell infiltration in iWAT of young and aged *Adh5*^BKO^ mice compared with aged-matched controls (Fig. 3D). Together, these data imply that ADH5*-*deficient thermogenic adipocytes accelerate “inflammaging.”

**Figure 3.**
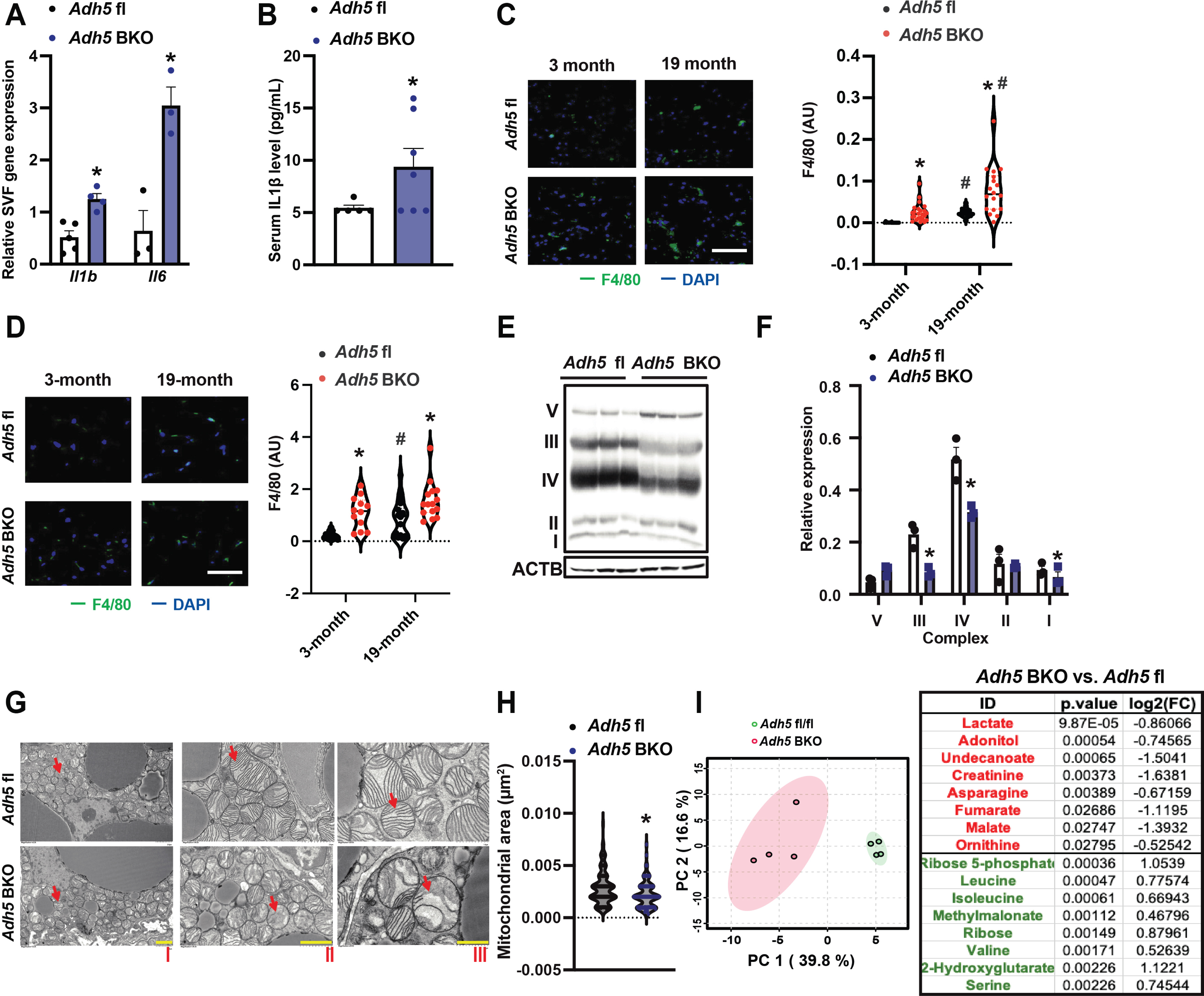
BAT ADH5*-*deficiency augments age-associated BAT immuno-metabolic imbalances. **A.** Gene expression of measured markers in SVF from BAT of *Adh5*^fl^ or *Adh5*^BKO^ mice. N= 3-5 mice/group**. B.** Plasma levels of IL-1β in 3-month-old *Adh5*^fl^ or *Adh5*^BKO^ mice measured by ELISA. N = 5-6 mice/group**. C.** Representative images (63X) and quantification of IF of F4/80 staining in BAT from *Adh5*^fl^ or *Adh5*^BKO^ mice at indicated age. N= 3 mice/group, 4-8 fields/mouse. Scale bar: 1 μm**. D.** F4/80 (scale bar: 1 μm) staining and quantification in iWAT from *Adh5*^fl^ or *Adh5*^BKO^ mice at the indicated age. N= 3 mice/group, 4-6 fields/mouse. **E&F.** Representative Western blot and quantification of ETC proteins in BAT of 3-month-old *Adh5*^fl^ or *Adh5*^BKO^ mice (n= 3 mice/group). **G**. Representative TEM images of BAT from 3-month-old *Adh5*^fl^ or *Adh5*^BKO^ mice. Red arrow, the mitochondrion. Scale bar: 5 μm (I), 2 μm (II), and 1 μm (III). **H.** Quantification of mitochondrial area in TEM images of BAT in mice in (C). Quantification was performed using ImageJ software**. I.** BAT metabolomic analysis in *Adh5*^fl^ or *Adh5*^BKO^ mice (3-month-old) analyzed by using Metaboanalyst online software; n = 4 age-matched mice per group. Left panel is the principal component analysis (PCA) and the right table is the list of the top eight metabolites downregulated (depicted in red) and upregulated (depicted in green) by ADH5 deletion. Data are presented as means ± SEM. * indicates genetic effects in the same age mice, and # indicates age effects in the same type of mice as determined by Student’s *t*-test (A, B, F and H), and two-way ANOVA (C&D), p<0.05.

Previously, we demonstrated that BAT-ADH5 deletion impairs BAT mitochondrial respiration in mice in response to cold and high-fat diet challenges[40]. Here, we found that BAT-ADH5-deletion decreased expression of the oxidative phosphorylation complexes I, III, and IV- key components of the mitochondrial electron transport chain (ETC) (Fig. 3E&F). Further assessment using transmission electron microscopy (TEM) analysis revealed smaller and aberrant cristae morphologies in BAT of *Adh5*^BKO^ mice compared to controls (Fig. 3G&H). To determine the impact of the loss of ADH5 on BAT metabolic homeostasis, we next carried out a steady-state metabolomic analysis in BAT from age-matched *Adh5*^BKO^ and *Adh5*^fl^ mice using gas chromatography-mass spectrometry (GC-MS). As shown in Fig. 3I, deletion of BAT ADH5 significantly decreased glycolytic metabolites and TCA cycle intermediates, and elevated levels of branched-chain amino acids (BCAAs) in BAT of young mice. Together, these data implicate that BAT ADH5 is required to maintain mitochondrial morphology and metabolic function.

### 3.4 Activation of BAT-HSF1 improves aging-associated outcomes

HSF1 is the master regulator of the heat shock response (HSR), a highly conserved physiological stress response to diverse stimuli[64]. We previously showed obesity suppresses BAT HSF1 and the HSF1-induced ADH5 expression[40]. Here we found aging downregulates BAT HSF1 at both transcript and protein levels starting from 6 months of age in male mice but not in female mice (Fig. 4A&B). In addition, we do not find significant effect of age on *Hsf1* expression in iWAT (Fig. 4C). Chromatin immunoprecipitation (ChIP) assay further confirmed that HSF1 binds to the *Adh5* promoter which was increased by a pharmacological HSF1 activator HSF1A (Fig. 4D)[40]. Consistently, pharmacological activation (HSF1A) or inhibition (KRIBB11) of HSF1 in BAT explants from wild-type mice significantly increased or decreased the BAT *Adh5*, respectively (Fig. 4E).

**Figure 4.**
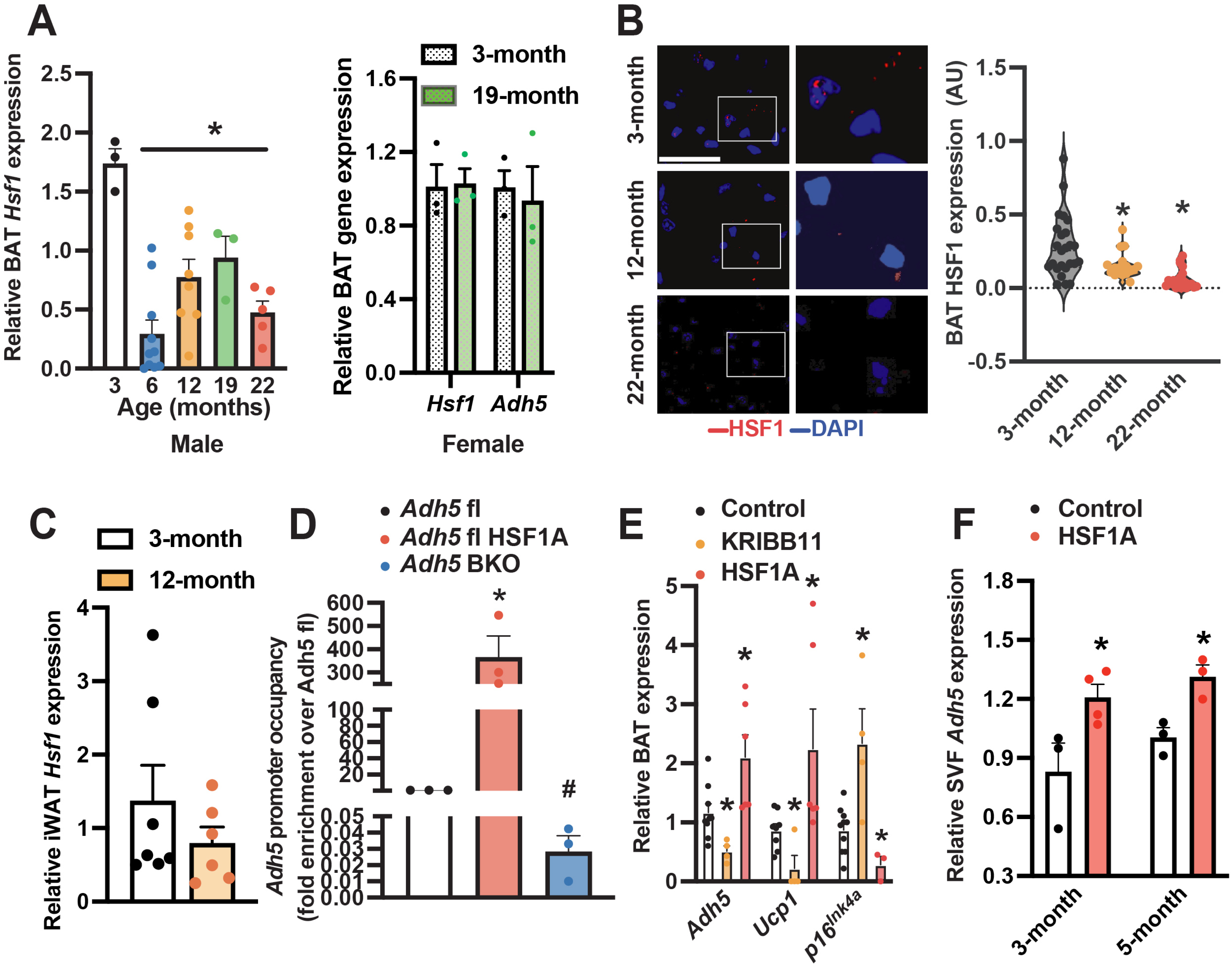
HSF1 induces *Adh5* and ameliorates senescence in BAT. **A.** *Hsf1* levels in BAT of male (n= 6-12 mice/group) and *Hsf1* and *Adh5* levels in BAT of female mice (n= 3 mice/group) at indicated age measured by qRT-PCR. Data were normalized to *Gapdh.* **B**. Left panel: representative HSF1 staining (63X, scale bar: 1 μm; zoomed images are on the right) from BAT of young and old wild-type mice. Right panel: quantification, n = 3-4 mice/age. 4-8 fields/mouse. **C.** *Hsf1 levels* in iWAT of male mice (n= 3 mice/group) at indicated age measured by qRT-PCR. Data were normalized to *Gapdh.* **D.** Occupancy of *Adh5* promoter regions by HSF1 from BAT treated with a pharmacological activator of HSF1 (HSF1A; Millipore, 20 μM, 24-hr) in controls or *Adh5*^BKO^ mice (n= 3 mice/group). **E**. Levels of mRNAs encoding the indicated genes of BAT explants treated ex vivo with HSF1A (20 μM, 24-hr) or HSF1 inhibitor (KRIBB11; 2 μm, 24-hr). Data were normalized to *Gapdh.* N=4 mice/group. **F.** Levels of mRNAs of the indicated genes in SVF from 3- or 5-month-old BAT pad isolated from wild-type mice followed by treatment of HSF1A (20 μM, 24-hr). N=3-4 mice/age group. Data were normalized to *Gapdh.* Data are presented as means ± SEM. *indicates statistical significance compared to the 3-month groups (A&B&C), to the *Adh5*^fl^ groups in (D), and to the control groups in (E-F) as determined by ANOVA in (A-male, B, and E), and Student’s *t*-test in (A-female, C, and F), p<0.05.

To establish the functional impact of the BAT HSF1-ADH5 axis in the context of aging, we utilized a nano clay (Nanosilicates, NS/NC)-mediated drug carrier system to ensure optimal drug-releasing efficiency, biocompatibility, and biodegradability[65–69]. Nanoclay mineral materials have excellent physicochemical characteristics necessary for drug delivery[70]. Here, collagen was used as a scaffold for NC-mediated HSF1A or vehicle delivery. As shown in Fig. 5A, compared with the collagen-HSF1A group, collagen-NC-HSF1A was effective in sustainably releasing compounds. Therefore, the collagen-NC-vehicle or collagen-NC-HSF1A formulation was injected interscapularly into the BAT of aged mice [40]. We found that one month after injection, activation of HSF1 significantly improved age-related systemic energy imbalance (Fig. 5B&C) and glucose intolerance (Fig. 5D). Moreover, these metabolic improvements were correlated with mitigated age-associated cognitive dysfunction (Fig. 5E&F), BAT morphology, as well as expression of senescent and inflammatory markers (Fig. 5G-J). Together, these data support the physiological relevance of BAT-HSF1 signaling in ameliorating aging-associated functional decline.

**Figure 5.**
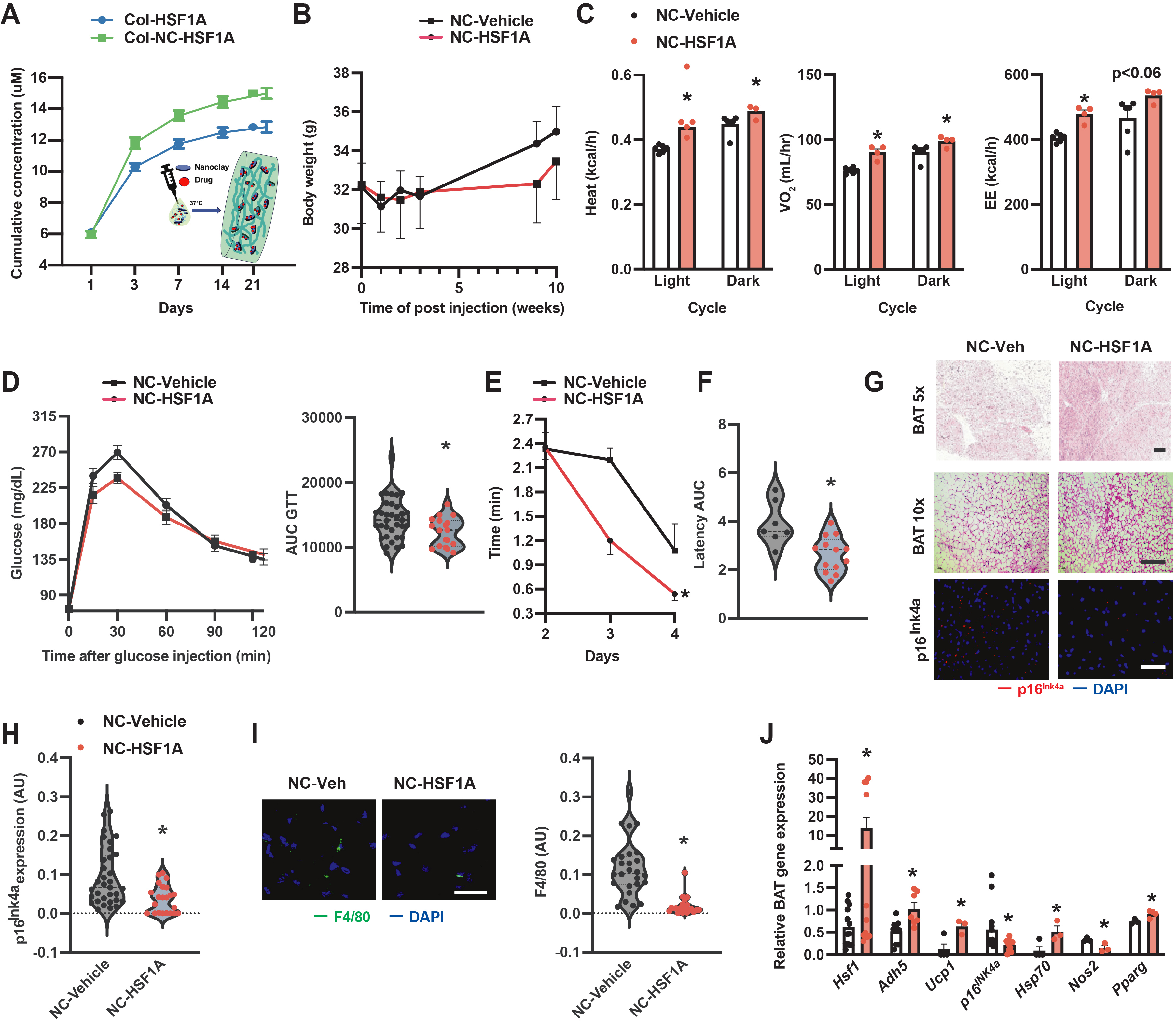
BAT HSF1 mitigates age-associated functional decline. **A.** Comparison of release curves between Collogen-NC-HSF1A vs Collagen-HSF1A. The amount of released HSF1A was quantified by UV–Vis spectroscopy using a standard curve for HSF1A. **B**. Body weight measurement over time in 12-month-old wild-type mice after interscapular injection with Collagen-NC-Vehicle (NC-Vehicle) or Collagen-NC-HSF1A (NC-HSF1A; 0.1 mg). **C.** Heat, VO_2,_ and energy expenditure (EE) in 12-month-old wild-type mice after one BAT injection of control (NC-Vehicle) or NC-HSF1A (0.1 mg; n= 5-6 mice/group). **D**. GTT and AUC of mice from (B). N= 19-37 mice/group. **E.** Latency (Barnes maze test) and **F.** AUC of escape measured from mice in (C) after one-month injection. N= 7-13 mice/group. **G.** Representative H&E (Scale bar: 20 or 10 μm) and IF of p16^lNK4a^ (scale bar: 10 μm) in BAT from mice in (C). **H.** Quantitation of IF staining of p16^lNK4a^. N= 5 mice/group. 4-6 fields/images. **I.** Representative F4/80 staining of BAT from (C); quantitation is left of the image. Scale bar: 1 μm, n = 5 mice/group. 4-6 fields/images. **J.** Levels of mRNAs encoding genes of interest (normalized to *Gapdh*) in BAT from mice in (C). N = 3-9 mice/group. Data are presented as means ± SEM. *indicates statistical significance compared to the vehicle groups as determined by Student’s *t*-test, p<0.05.

## 4. DISCUSSION

Aging is associated with a decline in thermogenic adipose tissue function and mass, contributing to disruption of energy and metabolic homeostasis as well as the prevalence of metabolic disease. Currently, several mechanisms have been implicated in age-dependent BAT decline including mitochondrial dysfunction, aberrant proteostasis, inflammation, and defective beiging[34; 71]. Here we provide the first evidence and mechanism by which nitrosative signaling links proteostasis to metabolic function in thermogenic adipose tissues in the context of aging.

In humans, aberrant nitric oxide bioactivity has been implicated in a diverse set of physiologies including aging[36; 37; 72]. It has been demonstrated that aging significantly changes BAT intracellular glutathione balance and systemic plasma nitrogen/arginine signatures[73], all of which are key components of the nitrosative signaling cascade. Notably, in humans, around 60% of plasma proteins significantly associated with aging are S-nitrosylated[74]. Specifically, in the BAT, more than 15% of protein cysteines are subject to oxidation modulating BAT physiology[28; 75]. However, the functional significance of BAT nitrosative signaling in aging remains undefined. Here we demonstrate an inverse correlation between ADH5 and aging-induced general BAT protein S-nitrosylation which is in line with previous studies demonstrating the relevance of ADH5 in aging. For example, recent studies demonstrated that AHD5 plays a critical role in DNA damage repair and chromatin architecture, and loss of this regulation leads to multisystem disorders and cancer progression[76–79]. Additionally, a genome-wide analysis has linked an *ADH5*-encompassing gene cluster to human longevity[80]. Finally, alterations in lifespan negatively correlate with *Adh5* in senescent mouse embryonic fibroblasts, neurons, and peripheral blood mononuclear cells from aged humans[21]. We found that defective ADH5-regulated S-nitrosylation signaling accelerates BAT senescence. Future studies are needed to delineate the direct actions of ADH5 on the cellular senescence network in the BAT.

Previously, it was demonstrated that mice with germline ADH5 deletion display accelerated aging phenotypes including reduced immune responses, increased morbidity upon endotoxic shock, insulin resistance, skeletal muscle wasting, and accumulation of protein aggregates in the brain cortex which are associated with locomotor deficits[21; 72]. Systemically, we showed that mice with BAT ADH5 deletion displayed increased body weight gain, fat mass, and inflammation in aged mice compared with age-matched controls. These augmented aging-developed metabolic defects were concomitant with worsened sensorimotor and cognitive function. These results are in line with previous studies showing that dysregulation of ADH5-mediated protein S-nitrosylation compromises mitochondrial function in the heart[81] and brain[21] as well as mitochondrial dynamics in the context of aging[21]. Moreover, at the cellular level, we found that BAT ADH5 deficiency reduced ETC complex expression and disrupted mitochondrial morphology, which was associated with elevated BAT-branched chain amino acid (BCAA) levels implicating a potential defect in BAT fuel utilization in *Adh5*^BKO^ mice. It is demonstrated that lactate and BCAA-derived metabolites control BAT mitochondrial redox homeostasis[82; 83] which in turn plays a key role in maintaining BAT metabolic function[84; 85]. Interestingly, recent studies in humans implicate an inverse correlation between BCAAs and longevity[86]. Furthermore, in rodents, impaired BCAA catabolism in adipose tissues promotes age-associated metabolic defects[87; 88]. Finally, in both human and animal studies, aberrant plasma levels of BCAAs serve as biomarkers for neurodegenerative diseases such as Parkinson’s and Alzheimer’s[88–91]. It is of great interest to explore to what extent BAT ADH5 regulates BCAA utilization in the future.

Excessive NO production by parenchymal cells drives immune cell infiltration and polarization in aging-associated diseases[92–94] and obesity[95]. Moreover, aberrant NO bioactivity and nitrosative stress modulate transcription factor activities that directly regulate inflammatory and adipogenic signatures[94; 96–99]. We found that loss of *Adh5* in BAT elevates NO production from BAs[40] and increases proinflammatory genes in BAs and SVF (Fig. 3) suggesting that the excess NO released from *Adh5* deficient BAs might serve as the signal-inducing proinflammatory mediator expression and immune cell activation leading to impaired BA mitochondrial function.

It is well recognized that heat shock factor 1 (HSF1) expression and activity decline in senescent human cells, thereby potentiating proteostasis collapse and the aging process[100; 101]. Indeed, HSF1-deficient organisms develop aging-related neurodegenerative disorders mainly caused by the accumulation of aggregates from misfolded proteins[26; 102; 103]. However, the mechanisms underlying multiple abnormalities, including developmental defects and increased expression of inflammatory cytokines, are unclear. In whole-body HSF1 KO mice, HSF1 deletion leads to cold intolerance and disrupted brown and beige function[104]. Notably, HSF1 levels are more abundant in thermogenic adipocytes[105] and restoration of HSF1 in BAT protects against obesity-associated BAT dysfunction[40; 104; 106]. Here, we showed that the age-dependent impairment of HSF1-*Adh5* signaling cascades leads to disruption of BAT metabolic homeostasis. In contrast, activation of HSF1 using a novel nanoclay-mediated drug delivery system ameliorated age-associated cellular and systemic declines. Although the cause factors driving the age-suppressed HSF1 remain unknown and are of great interest for future studies, our study demonstrates a novel strategy to combat the thermogenic adipose decline in the context of aging. We recognize that nitrosative signals broadly influence cellular signaling cascades, and the functions of BAT are governed by diverse transcriptional and signaling programs, neuronal tone, as well as endocrine factors[1; 107–109]. Thus, we cannot rule out the possibility that the impact of *Adh5* on mitochondrial dynamics and fuel utilization in the BAT is secondary to the disruption of other cellular functions. In future studies, we will examine the effect of *Adh5* deletion on other organelle functions such as the endoplasmic reticulum and lysosomes.

Taken together, we provide the first evidence of NO bioactivity-modulated BAT immune-metabolic homeostasis as well as new mechanisms by which nitrosative signaling links proteostasis to metabolic function in thermogenic adipose tissue. Moreover, we highlight a new geroprotective strategy to counteract aging-associated cellular redox imbalance. Ultimately, this study will provide new avenues for developing novel therapeutic strategies to mitigate age-dependent loss of thermogenic adipocyte function and promote healthy aging.

## Author Contributions

S.C.S., T.N., V.A.L., H.S., M.J.P., and L.Y. designed the research; S.C.S., T.N., Q.Q., A.A., Z.W., M.L. and A.B. performed experiments; S.C.S., T.N., and L.Y. analyzed the data; and S.C.S., T.N., and L.Y. wrote the paper. S.C.S. and T.N. equal contribution. S.C.S., M.J.P., and L.Y. are supported by R01DK126817, and T.N. was supported by NIH 5T90DE023520.

## DUALITY OF INTEREST

No potential conflicts of interest relevant to this article were reported.

## ACKNOWLEDGMENTS

We are grateful to all Yang laboratory members for supporting this project.

## Notes

### Competing Interest Statement

The authors have declared no competing interest.

### Summary of Updates

We have included the data in Figure 3I instead of the schematic summary of metabolomics analysis.

